# Effects of Antibiotic Interaction on Antimicrobial Resistance Development in Wastewater

**DOI:** 10.1101/2023.02.10.528009

**Authors:** Indorica Sutradhar, Carly Ching, Darash Desai, Zachary Heins, Ahmad S. Khalil, Muhammad H. Zaman

## Abstract

While wastewater is understood to be a critically important reservoir of antimicrobial resistance due to the presence of multiple antibiotic residues from industrial and agricultural runoff, there is little known about the effects of antibiotic interactions in the wastewater on the development of resistance. We worked to fill this gap in quantitative understanding of antibiotic interaction in constant flow environments by experimentally monitoring *E. coli* populations under subinhibitory concentrations of combinations of antibiotics with synergistic, antagonistic, and additive interactions. We then used these results to expand our previously developed computational model to account for the complex effects of antibiotic interaction. We found that while *E. coli* populations grown in additively interacting antibiotic combinations grew predictably according to the previously developed model, those populations grown under synergistic and antagonistic antibiotic conditions exhibited significant differences from predicted behavior. *E. coli* populations grown in the condition with synergistically interacting antibiotics developed less resistance than predicted, indicating that synergistic antibiotics may have a suppressive effect on antimicrobial resistance development. Furthermore *E. coli* populations grown in the condition with antagonistically interacting antibiotics showed an antibiotic ratio-dependent development of resistance, suggesting that not only antibiotic interaction, but relative concentration is important in predicting resistance development. These results provide critical insight for quantitatively understanding the effects of antibiotic interactions in wastewater and provide a basis for future studies in modelling resistance in these environments.

**Importance:** Antimicrobial resistance (AMR) is a growing global threat to public health expected to impact 10 million people by 2050, driving mortality rates globally and with a disproportionate effect on low- and middle-income countries. Communities in proximity to wastewater settings and environmentally contaminated surroundings are at particular risk due to resistance stemming from antibiotic residues from industrial and agricultural runoff. Currently, there is a limited quantitative and mechanistic understanding of the evolution of AMR in response to multiple interacting antibiotic residues in constant flow environments. Using an integrated computational and experimental methods, we find that interactions between antibiotic residues significantly affect the development of resistant bacterial populations.

## Introduction

Antimicrobial resistance (AMR) is a rapidly evolving critical threat to global health with the potential to lead to financial losses of as much as $100 trillion USD (1, 2). A recent systematic analysis of global AMR has predicted that there were an estimated 4.95 million deaths associated with bacterial AMR in 2019 (3). Contributing factors to AMR in human medicine (i.e., prescription patterns, poor patient treatment adherence etc.) have been well documented (4–7); however, environmental distribution of antibiotics and its impact on AMR has received less attention (8, 9). Wastewater specifically has been shown to be a reservoir of resistant pathogens, often stemming from the antibiotic pollution present in runoff from industrial and agricultural sources (10). Furthermore, computational modeling of wastewater has shown that even low concentrations of antibiotic residues can lead to the development of AMR (11). This is particularly of concern in low-income communities which can often have open sewer systems and little access to wastewater treatment, putting them at particular risk for deadly drug-resistant outbreaks.

Previously, we have developed a computational model of resistance acquisition in continuous flow environments based on known mechanisms of bacterial growth and mutation as well as experimental validation (11). However, experimental validation of the model was limited to systems with only one antibiotic residue. The interaction between two or more antibiotics is of particular interest, with combination therapy used both clinically to increase treatment efficacy and lower the risk of AMR development as well as prophylactically in livestock to prevent infections from developing and spreading across these large animal populations. The interaction between two antibiotics from different classes have previously been shown to affect resistance acquisition (12). Synergy is the interaction of multiple drugs to have a greater killing action than the sum of their parts while antagonism is the interaction of multiple drugs to have reduced killing action than the sum of their parts. Drugs that do not interact, or in other words have the killing action equal to the sum of their parts are said to have an additive interaction. Interestingly, synergy between two antibiotics has also been shown to increase the likelihood of resistance population development at subtherapeutic doses (12). However, the effects of antibiotic interaction on the growth of resistant populations in wastewater settings has not previously been observed. Wastewater can often have many antibiotic residues present, which have the potential to interact with each other either synergistically or antagonistically. For example, antibiotic residues found in water sampled from hospital sewage in Sweden included the drugs doxycycline, erythromycin, and ciprofloxacin among others (13). This is of note because doxycycline and erythromycin are known to have a synergistic interaction, while doxycycline and ciprofloxacin are known to have an antagonistic interaction (12). Antimicrobials in combination often have different mechanisms of action, so it is possible that interactions between the multiple antibiotic residues in wastewater will have unique effects on the development of antimicrobial resistance. However, quantitative data on these effects of antibiotic interactions on AMR in wastewater is lacking. We aim to fill this critical gap in knowledge about the effects of antibiotic interactions on AMR in continuous flow environments such as wastewater through an iterative approach to computational modeling and experimental validation.

## Methods

### Model Development

The model used in this paper is based on a previously developed model of the growth of antibiotic resistant bacterial populations in wastewater that builds on prior studies and extends to incorporate a variety of critical inputs which can be broadly classified into bacterial parameters, environmental parameters and antibiotic parameters (11). Bacteria specific input factors include the growth rates of antibiotic susceptible and resistant strains and mutation rates in response to subinhibitory concentrations of antibiotic. The antibiotic specific inputs, such as bactericidal activity, allow for the study of the effects of antibiotic pollution on the development of resistance. Additionally, environmental inputs, including physical inflow and outflow rates and antibiotic residue concentrations, allow for the modelling of resistance development in a variety of settings of interest. Ordinary differential equations incorporating these input parameters were used to model an output of resistant bacterial populations over time, thus allowing for the prediction of resistant population development (Eq Set 1 and Table 1).

**Table 1.** Model variables and definitions

| Variable | Definitions |
| --- | --- |
| $C_1$ | Antibiotic 1 Concentration (ug/mL) |
| $C_2$ | Antibiotic 2 Concentration (ug/mL) |
| $S$ | Susceptible (cells) |
| $R_m$ | Resistant to both Antibiotic 1 and Antibiotic 2 from Chromosomal Mutation (cells) |
| $R_1$ | Resistant to only Antibiotic 1 from Chromosomal Mutation (cells) |
| $R_2$ | Resistant to only Antibiotic 2 from Chromosomal Mutation (cells) |
| $E$ | Environmental Concentration of Antibiotic((ug/mL)/hr) |
| $syn$ | Synergy Parameter (non-dimensional) |
| $k_e$ | Antibiotic Clearance (1/hr) |
| $\alpha_S$ | Growth Rate of Susceptible Bacteria (1/hr) |
| $\alpha_{Rm}$ | Growth Rate of Bacteria Resistant from Mutation (1/hr) |
| $N_{max}$ | Carrying Capacity (cells/mL) |
| $g_S, g_{Rm},$<br>$g_{R1}, g_{R2}, g_{Rp}$ | Bacterial Influx Rates (cells/hr) |
| $k_T$ | Bacterial Efflux Rate (1/hr) |
| $\delta_{max,1}, \delta_{max,2}$ | Bacterial Killing Rate in Response to Antibiotic 1 and Antibiotic 2 (1/hr) |
| $C_S^{50}, C_{R,1}^{50},$<br>$C_{R,2}^{50}$ | Antibiotic Concentration where the Killing Action is Half its Maximum Value (ug/mL) |
| $m_1(C_1)$ | Mutation Frequency under Antibiotic 1 (1/hr) |
| $m_2(C_2)$ | Mutation Frequency under Antibiotic 2 (1/hr) |

### Experimental Validation

Experimental validation of the model was done using the eVOLVER system, which is an automated, highly flexible platform allowing for scalable continuous culture microbial growth and independent, precise and multiparameter control of growth conditions such as temperature and flow rate (14). Experiments were done with antibiotics which have been found to be present in wastewater with known interactions with one pair of antibiotics exhibiting additive interaction (12.5 mg/L Rifampicin + 4 mg/L Streptomycin), one pair of antibiotics exhibiting synergistic interaction (1.5 mg/L Doxycycline + 64 mg/L Erythromycin) and one pair of antibiotics exhibiting antagonistic interaction (1.5 mg/L Doxycycline + 0.0375 mg/L Ciprofloxacin) (12-13, 15-17). Drug interactions were confirmed using checkerboard assays and calculating fractional inhibitory concentration (FIC) values as described in Bellio et al. where combinations with an FIC less than 0.5 were considered synergistic, those with an FIC greater than 4 were considered antagonistic and those with an FIC between 0.5 and 4 were considered to have an additive interaction (18). Experiments were initialized with inoculation of LB media with *E. coli* MG1655 in static conditions at 37°C. Then, inflow and outflow of the antibiotic-containing LB media at two concentration combinations was started. During the course of the experiment, each culture condition was sampled daily, and the concentrations of total bacteria and resistant bacteria were calculated through plating on drug-free and selective LB agar containing 8X MIC Drug A and/or 8X MIC Drug B respectively.

## Results

### Drugs with Additive Interaction Develop Resistance Predictably

The first antibiotic combination tested was Rifampicin and Streptomycin at half of their respective MICs (12.5 mg/L Rifampicin + 4 mg/L Streptomycin). Checkerboard assays confirmed an FIC of 1 indicating additive interaction between these two drugs. Model prediction was made based on previously determined parameter values from eVOLVER experiments with each drug in isolation. The experimental results with the eVOLVER qualitatively verified the model prediction with dominant susceptible and Rifampicin-resistant populations as well as a significant population of bacteria resistant to both Rifampicin and Streptomycin (Figure 1).

**Figure 1:**
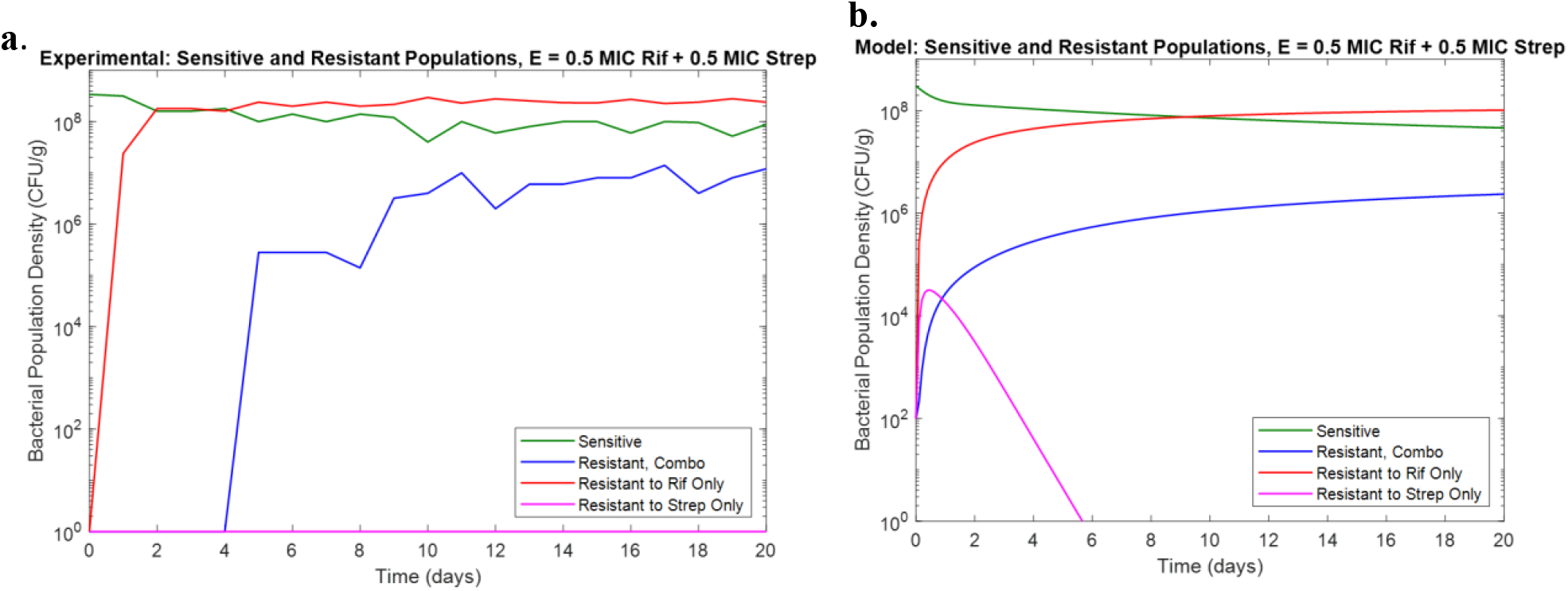
**a.)** Experimental results for combination of 0.5X MIC Rifampicin and 0.5X MIC Streptomycin in eVOLVER. **b.)** Model prediction for combination of 0.5X MIC Rifampicin and 0.5X MIC Streptomycin in eVOLVER

While a population of bacteria resistant to Streptomycin only was not observed experimentally, this may be due to the transient nature of this population not being captured in the sampling frequency. This confirmed the assumption that antibiotics with no interaction behave predictably in combination.

### Synergistic Interaction Show Lower than Expected Resistance

The second antibiotic combination tested was Doxycycline and Erythromycin, which in addition to have been observed in wastewater sampling, have also previously found to interact synergistically (13, 17). Checkerboard assays confirmed an FIC of 0.375 indicating synergistic interaction. Initial model prediction was made based on previously determined parameter values from eVOLVER experiments with each drug in isolation and assuming no effect from antibiotic interaction, showing dominant Doxycycline resistant and combination resistant populations (Figure 2a). Experimental results showed lower levels of resistance than predicted, particularly in the bacterial population resistant to both drugs (Figure 2b). In order to reproduce the experimental behavior, a synergy parameter, equal to the FIC value for the given antibiotic combination, was then introduced as a multiplying factor to the mutation parameter to account for reduced resistance levels (Eq set 1). The results of this change are shown in Figure 2c. These results suggest that synergy may have a suppressive effect on the development of resistance due to a decrease in the mutation rates proportional to the degree of synergy. This is of particular interest because previous studies done in non-flow conditions saw increased resistance in synergistic conditions compared to antibiotics with no interaction, indicating that environments with constant flow cannot be adequately predicted with only data from standard non-flow culture conditions (17).

**Figure 2:**
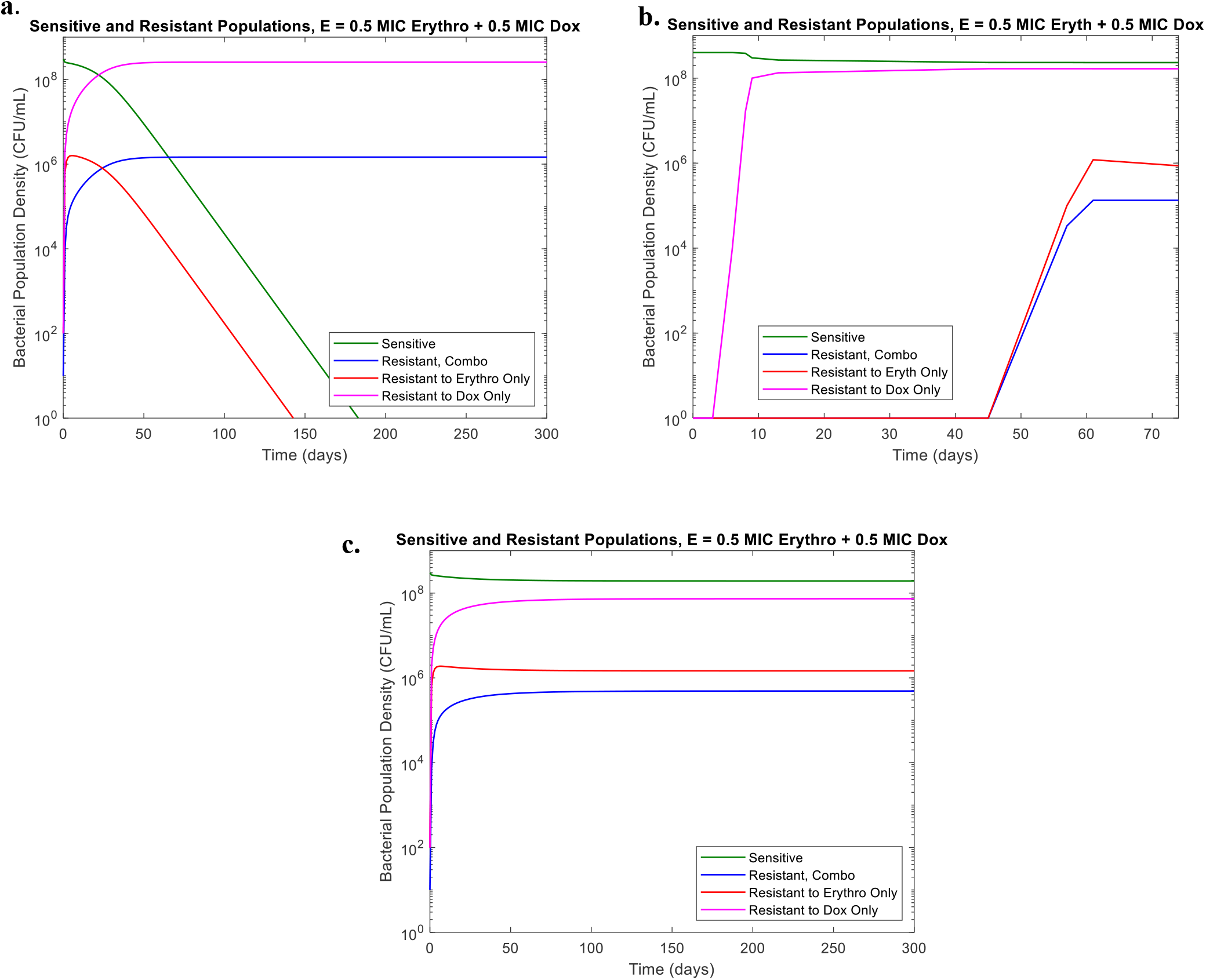
**a.)** Model prediction for combination of 0.5X MIC Doxycycline and 0.5X MIC Erythromycin in eVOLVER in absence of interaction **b.)** eVOLVER results for combination of MIC Doxycycline and 0.5 MIC Erythromycin **c.)** Model prediction for combination of 0.5X MIC Doxycycline and 0.5X MIC Erythromycin in eVOLVER with synergy parameter

### Drugs with Antagonistic Interaction Exhibit Ratio-Dependent Resistance Development

The third antibiotic combination tested was Doxycycline and Ciprofloxacin which have been observed as residues in wastewater samples and have previously found to interact antagonistically (13, 17). Checkerboard assays confirmed an FIC of 4 indicating antagonistic interaction. Initial experimental results showed lower levels of resistance than predicted, and no observable bacterial population resistant to both drugs (Figure 3a). However, previous studies indicated that unlike in additive and synergistic combinations, resistance development in Doxycycline and Ciprofloxacin combinations may differ depending on the relative concentrations of the two (17). Additional experiments were conducted with differing ratios of Doxycycline and Ciprofloxacin (0.9 MIC Dox: 0.1 MIC Cip; 0.7 MIC Dox: 0.3 MIC Cip; 0.5 MIC Dox: 0.5 MIC Cip; 0.3 MIC Dox: 0.7 MIC Cip; 0.1 MIC Dox: 0.9 MIC Cip). These experiments found an antibiotic-ratio dependent effect. Several changes were made to the previous model to account for the ratio dependency of the resistant population behavior (Table 2). First, the growth term was adjusted to include an antibiotic concentration dependent growth rate function rather than a constant growth rate parameter. Additionally, the bacterial killing rate parameter was similarly adjusted to include a resistant population-dependent killing rate function rather than a constant killing rate. The results of the adjusted model for the 50% MIC Dox and 50% MIC Cip condition are shown in Figure 3b, demonstrating the model’s ability to capture the dominant susceptible population as well as the lower Doxycycline resistant population and the absence of the combination resistant population as seen in Figure 3a.

**Figure 3:**
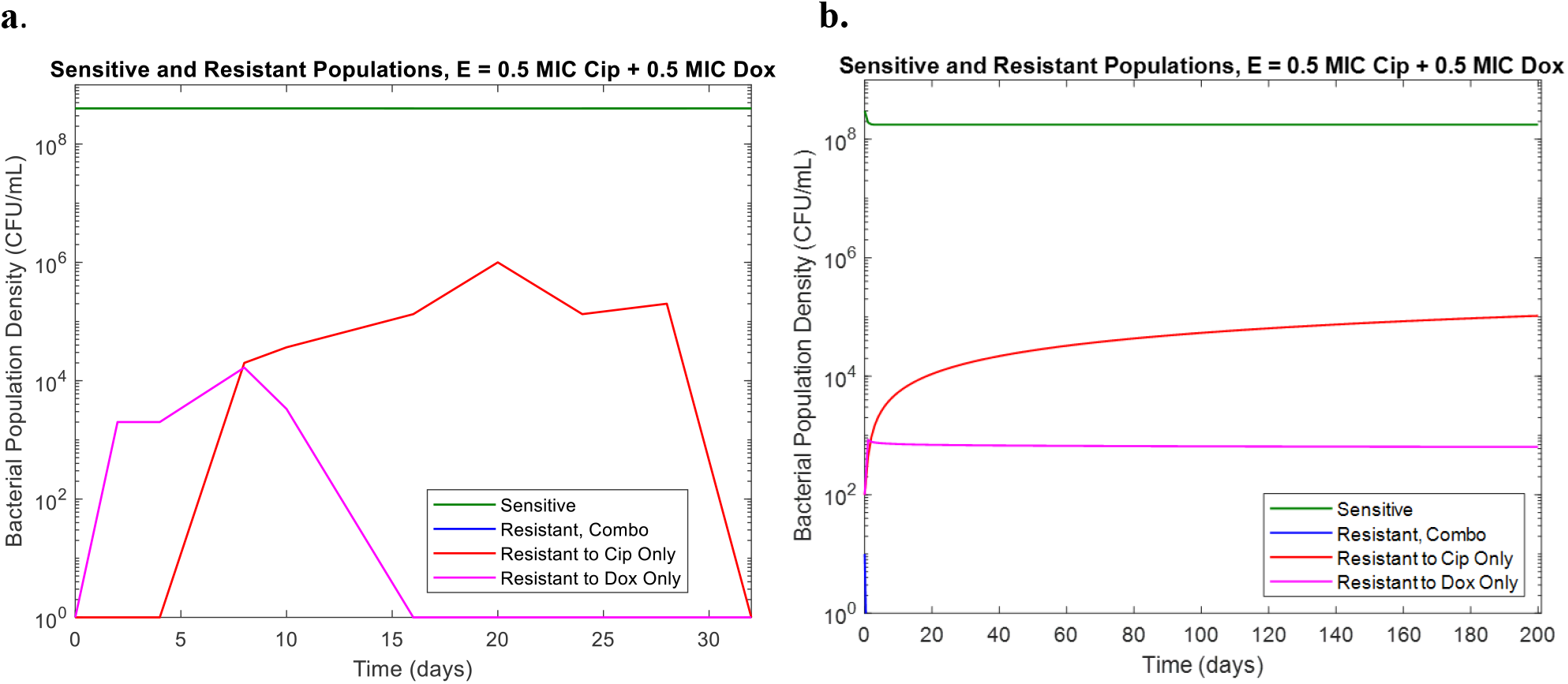
**a.)** eVOLVER results for combination of 0.5 MIC Doxycycline and 0.5 MIC Ciprofloxacin **b.)** Model prediction for combination of 0.5 MIC Doxycycline and 0.5 MIC Ciprofloxacin in eVOLVER with antibiotic ratio dependent model adjustments

**Table 2:** Model adjustment for antibiotic-ratio dependent antagonistic behavior

| Base Model | Adjusted Model for Antagonism |
| --- | --- |
| Growth Term:<br>$\alpha_{R,2} * \left(1 - \frac{R_m + R_1 + R_2 + S}{N_{max}}\right) R_2$ | Growth Term: $\alpha_{R,2}(MIC1, MIC2) * \left(1 - \frac{R_m + R_1 + R_2 + S}{N_{max}}\right) R_2$<br>Where : $\alpha_{R,2}(MIC1, MIC2) = \alpha_{R,2} * sup1 * \left(\frac{MIC1}{MIC2}\right)$ |
| Bacterial Killing from Dox:<br>$syn * \delta_{max,2} \left(\frac{C_2}{C_2 + C_{R,2}^{50}}\right) R_2$ | Bacterial Killing from Dox: $syn * \delta_{max,2}(R_1, R_2) * \left(\frac{C_2}{C_2 + C_{R,2}^{50}}\right) R_2$<br>Where: $\delta_{max,2}(R_1, R_2) = \delta_{max,2} * sup2 * \left(\frac{R_1}{R_2}\right)$ |

| Parameter | Description |
| --- | --- |
| $sup1$ | Cip Suppression Variable 1 |
| $sup2$ | Cip Suppression Variable 2 |
| $\alpha_S$ | Growth Rate of Susceptible Bacteria (1/hr) |
| $\alpha_R$,<br>$\alpha_{R,1}, \alpha_{R,2}$ | Growth Rate of Bacteria Resistant from Mutation (1/hr) |
| $N_{max}$ | Carrying Capacity (cells/mL) |
| $MIC1$ | MIC in response to Cip |
| $MIC2$ | MIC in response to Dox |
| $\delta_{max,1}$,<br>$\delta_{max,2}$ | Bacterial Killing Rate in Response to Antibiotic 1 and Antibiotic 2 (1/hr) |

This adjusted model was able to capture the relative behaviors of the different resistant populations for differing ratios of Dox and Cip, notably the transient Doxycycline-resistant population giving way to the combination resistant population (Figure 4). Furthermore, the model successfully captures the increased time the Doxycycline-resistant population was present in the condition with 0.9X MIC Dox (Figures 4c-d) compared to the condition with 0.7X MIC Dox (Figures 4a-b). However, it failed to capture the sustained drug-susceptible population in the high Cip concentration conditions. We hypothesize that this may be due to a separation of the drug susceptible populations from the resistant population between the planktonic bacteria and the bacteria in the biofilm that form at walls of the eVOLVER vials. Biofilm has been seen to have different resistance profiles than planktonic bacteria which may explain why the model, which only accounts for the bacteria under constant flow conditions, does not fully capture the susceptible population (19). Though the current model is limited in its ability to model bacteria in biofilm, it still succeeds in being able to predict the resistance development occurring in the continuous liquid culture. Thus, it still can have use as a predictive tool for understanding AMR in wastewater.

**Figure 4:**
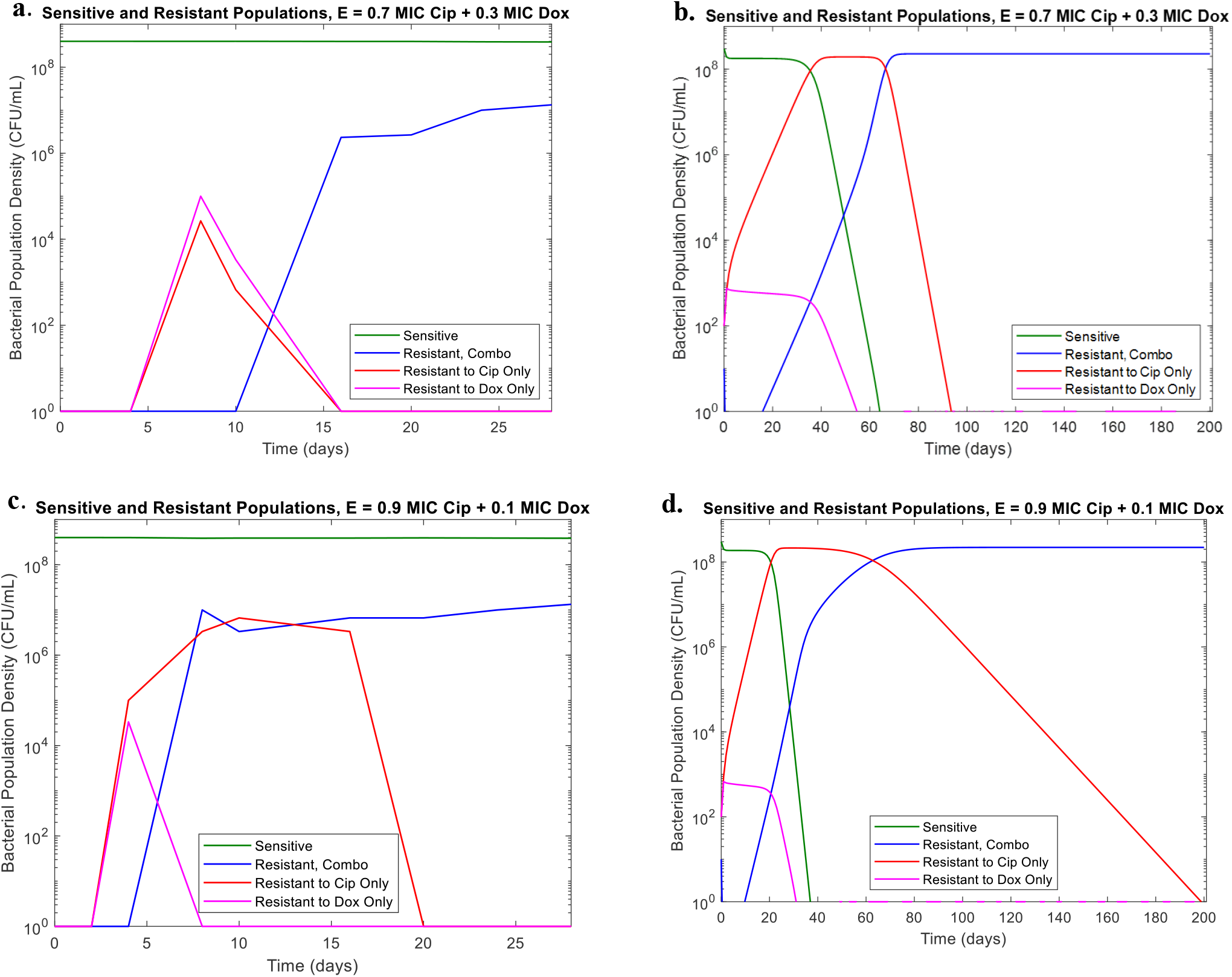
**a.)** eVOLVER results for combination of 0.3 MIC Doxycycline and 0.7 MIC Ciprofloxacin in eVOLVER in absence of interaction **b.)** Model prediction for combination of 0.3 MIC Doxycycline and 0.7 MIC Ciprofloxacin in eVOLVER with antibiotic ratio dependent model adjustments **c.)** eVOLVER results for combination of 0.1 MIC Doxycycline and 0.9 MIC Ciprofloxacin in eVOLVER in absence of interaction **d.)** Model prediction for combination of 0.1 MIC Doxycycline and 0.9 MIC Ciprofloxacin in eVOLVER with antibiotic ratio dependent model adjustments

## Discussion and Conclusions

Overall, through our integrated computational and experimental approach, we were able to model the development of antibiotic resistance in response to subinhibitory combinations of antibiotics exhibiting additive, synergistic and antagonistic interactions. We demonstrated *E. coli* populations grown in additively interacting antibiotic combinations grew predictably according to the previously developed model. This confirmed our assumption that in the absence of antibiotic interaction, resistance to each antibiotic will develop independently. We also found that *E. coli* populations grown under synergistic and antagonistic antibiotic conditions exhibited significant differences from predicted behavior. *E. coli* populations growing in subinhibitory concentrations of synergistically interacting antibiotics showed the development of less resistance than predicted. Interestingly, this indicated that synergistic antibiotics have a suppressive effect on antimicrobial resistance development in continuous flow conditions. This is in contrast to previous studies in non-flow conditions which found that synergy increased resistance acquisition (12, 20). Thus, our novel finding suggests that differing flow conditions significantly alter resistance acquisition patterns and studies in continuous flow conditions are necessary for understanding environments like wastewater. Additionally, we found that *E. coli* populations grown with antagonistically interacting antibiotics showed an antibiotic ratio-dependent development of resistance. This behavior has previously been observed in non-flow conditions, though only with single-resistant populations (17). Our studies further this finding to multi-resistant populations and also find that not only antibiotic interaction, but relative concentration is important in predicting resistance development in continuous flow environments.

Though we have been able to draw a number of conclusions about the effects of antibiotic interaction on resistance development in wastewater, we note that our studies do have limitations. Primarily, we only studied a limited number of antibiotic combinations and as such cannot conclude the effects of all antibiotic interactions. Future studies investigating a wider array of antibiotics could elucidate further findings on specific combinations in constant flow conditions. Additional studies looking at three or more antibiotics in combination with varying interactions or media conditions better approximating wastewater than the LB broth used here would also be a step forward in modelling the types of complex conditions that would be found in wastewater. Another major area of interest in developing the model would be to further integrate the role of biofilm in resistance development. While biofilm has been observed in samples both up- and downstream from wastewater treatment plants and is a known environmental reservoir of resistance, there is limited quantitative understanding of how this resistance develops, particularly in response to antibiotic residues present in wastewater (21, 22). In order to develop quantitative models of resistance development in wastewater incorporating both planktonic bacteria and biofilm, experimental methods for controllably maintaining both populations in continuous flow conditions will need to be developed.

Despite these limitations, experimental validation demonstrated our ability to model resistance development in subinhibitory antibiotic concentrations of antibiotics with varying interactions. We were able to determine that synergistic interaction have a suppressive effect on resistance development. Additionally, more complex resistance development patterns were observed in the case of antagonistic interaction where we found an antibiotic ratio-dependent behavior. This has important implications for understanding the effects of industrial and agricultural antibiotic runoff in wastewater and determining acceptable antibiotic concentrations and combinations when treating wastewater. These findings can be used as a basis for public health policy makers and the developed model can be utilized to predict resistant population emergence in different sewage and wastewater conditions where multiple antibiotic residues may be present.

## Author Contributions

I. Sutradhar designed the model, conducted experiments, and analyzed the data. C. Ching and D. Desai provided guidance on model design and verification. A. Khalil provided facilities for experimental work with the eVOLVER and Z. Heins provided assistance on experimental design. I. Sutradhar and M. H. Zaman wrote the article.

## Acknowledgements

This research was supported in part by the National Institutes of Health (NIH) training grant at Boston University, T32 EB006359 (to I. S.) and the NIH grant R01AI171100 to A.S.K. A.S.K. also acknowledges funding from a NIH Director’s New Innovator Award (DP2AI131083) and a Department of Defense Vannevar Bush Faculty Fellowship (N00014-20-1-2825). The content is solely the responsibility of the authors and does not necessarily represent the official views of the National Institutes of Health.

## Tables and Equations

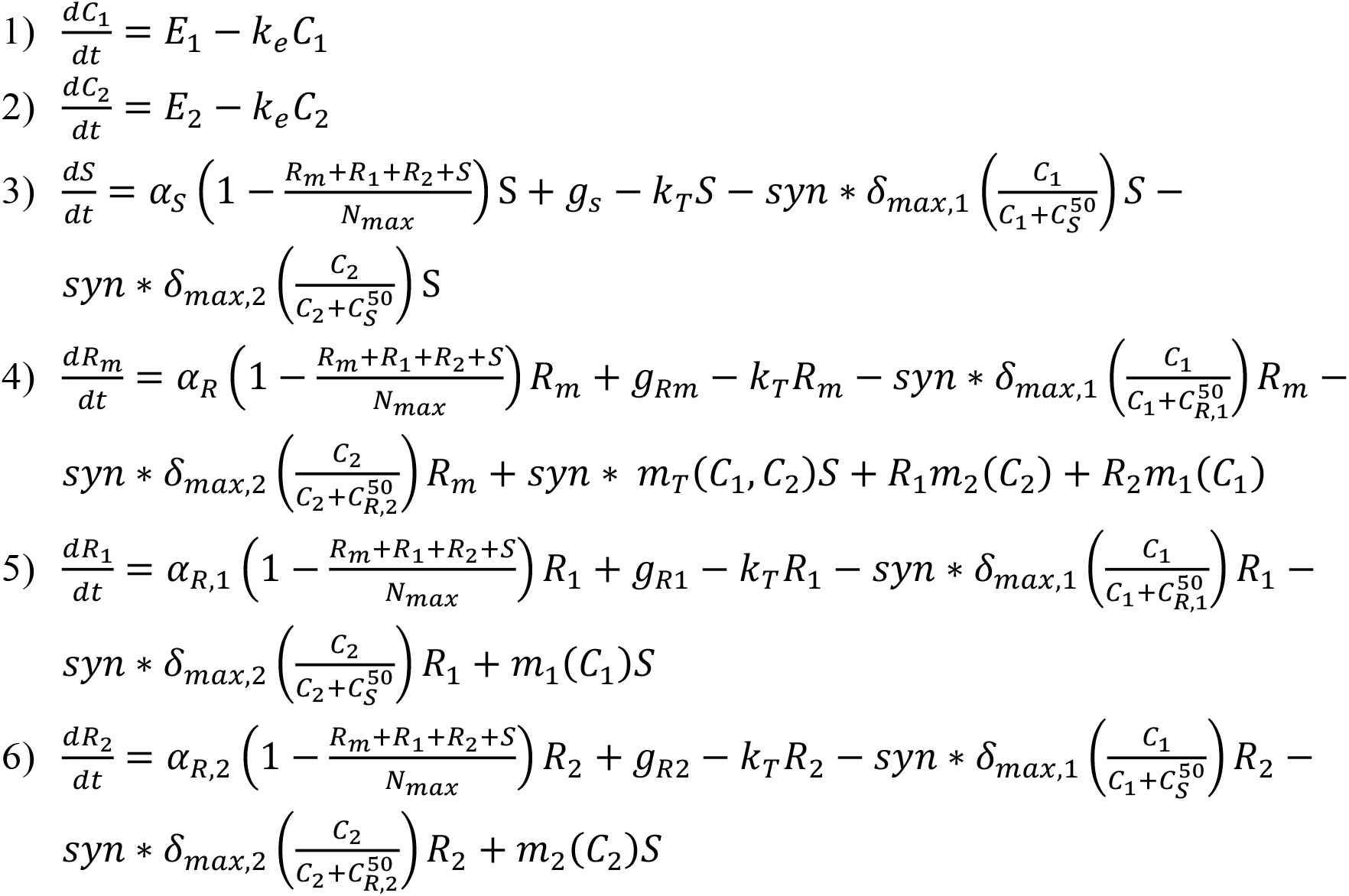

**Eq Set 1**. Sensitive and resistant populations under selective pressure from antimicrobial combination therapy, adapted from Sutradhar et al. 2021^11^

